# Cross-feeding modulates antibiotic tolerance in bacterial communities

**DOI:** 10.1101/243949

**Authors:** Elizabeth M. Adamowicz, Jeffrey Flynn, Ryan C. Hunter, William R. Harcombe

## Abstract

Microbes frequently rely on metabolites excreted by other bacterial species, but little is known about how this cross-feeding influences the effect of antibiotics. We hypothesized that when species rely on each other for essential metabolites, the minimum inhibitory concentration (MIC) for all species will drop to that of the “weakest link” - the species least resistant in monoculture. We tested this hypothesis in an obligate cross-feeding system that was engineered between *Escherichia coli, Salmonella enterica*, and *Methylobacterium extorquens*. The effect of tetracycline and ampicillin were tested on both liquid and solid media. In all cases, resistant species were inhibited at significantly lower antibiotic concentrations in the cross-feeding community than in monoculture or a competitive community. However, deviation from the “weakest link” hypothesis was also observed in cross-feeding communities apparently as result of changes in the timing of growth and cross-protection. Comparable results were also observed in a clinically relevant system involving facultative cross-feeding between *Pseudomonas aeruginosa* and an anaerobic consortium found in the lungs of cystic fibrosis patients. *P. aeruginosa* was inhibited by lower concentrations of ampicillin when cross-feeding than when grown in isolation. These results suggest that cross-feeding significantly alters tolerance to antibiotics in a variety of systems.

## Introduction

Antibiotic resistant bacteria pose a considerable public health threat worldwide; the World Health Organization reports that 25–50% of hospital-acquired pathogens are now multiple-drug-resistant (World Health Organization., 2014). Despite extensive research on the cellular mechanisms of resistance in many bacterial species (Friman et al., 2015; Hughes and Andersson, 2012), a growing body of research suggests that a single-species view of pathogen response to an antibiotic may be incomplete. Many infections are now known to involve multiple pathogens (de Vos et al., 2017; Tay et al., 2016) or interactions between pathogens and commensals, (Delhaes et al., 2012; Murray et al., 2014; Ramsey et al., 2011). As well, we still have little understanding of how ecological interactions between microbes influence the impact that antibiotics have on populations and communities of bacteria.

Growth in a polymicrobial consortium can influence a species’ antibiotic tolerance by multiple mechanisms (Perlin et al., 2009; Yurtsev et al., 2016). First, resistant species can protect more sensitive species by degrading antibiotics. For example, β-lactamase production leads to the degradation of β-lactam antibiotics, thereby detoxifying shared growth medium (Medaney et al., 2016; Perlin et al., 2009). Additionally, secretions from one species can induce resistance mechanisms in others. Secretion of the signalling molecule indole has been shown to increase resistance by activating stress-response pathways in *S. enterica* (Vega et al., 2013) and efflux pump expression in *Pseudomonas putida* (Molina-Santiago et al., 2014). Community growth induces physiological changes in bacteria (Straight and Kolter, 2009), which may alter their antibiotic tolerance by increasing drug uptake or slowing their metabolic rate (Hong et al., 2014; Peng et al., 2015). Further, secretions that change environmental pH have been suggested to alter the impact of antibiotics (Bernier et al., 2011; de Vos et al., 2017). Spatial structure may also play a role in these protective interactions; a synthetic community of *Pseudomonas aeruginosa, Pseudomonas protegens*, and *Klebsiella pneumoniae* was found to have greater tobramycin resistance when grown as a multi-species biofilm versus single species biofilm or multispecies planktonic culture (Lee et al., 2014). However, in many cases, the mechanisms underlying communities’ effects on resistance remain unclear (de Vos et al., 2017).

The primary mechanism of interaction between bacteria is through extracellular metabolites, yet few studies have investigated how metabolic interactions in a bacterial community modulate the impact of antibiotics (Manor et al., 2014). One such metabolic interaction, called cross-feeding, occurs when metabolites produced by one organism are used as a nutrient or energy source by another (Rosenzweig et al., 1994; Schink, 2002). This phenomenon is nearly ubiquitous in microbial communities (Pande et al., 2014; Seth and Taga, 2014; Zelezniak et al., 2015) and is thought to contribute to our inability to cultivate most bacterial species in isolation (Kaeberlein et al., 2002; Oliveira et al., 2014). Cross-feeding has also been shown to play a critical role in the human microbiome. For example, *Bifidobacterium* species in the gut cooperate with each other to break down complex plant-based carbohydrates (Milani et al., 2015), which releases carbon for butyrate-producing bacteria. Cross-feeding in the gut also increases the efficiency of glycan breakdown by removing waste products such as H_2_ (Koropatkin et al., 2012). Additionally, cross-feeding can influence pathogen survival. For example, growth and virulence of the oral pathogen *Aggregatibacter actinomycetemcomitans* depends on L-lactate generated by the commensal *Streptococcus gordonii* (Stacy et al., 2016). Given the ubiquity and importance of cross-feeding in human-associated microbial communities, greater investigation in to how cross-feeding influences population and community responses to antibiotics is needed.

In this work, we test how cross-feeding changes the effect of antibiotics on bacteria in microbial communities. We use the term tolerance to signify the ability of species to grow in a given antibiotic concentration. Tolerance as we define it can change as a function of physiological state or environmental conditions, while changes in resistance would require a change in DNA sequence (Brauner et al., 2016). We hypothesize that when species depend on one another the community tolerance (i.e. the level of antibiotic required to inhibit detectable growth of the community as a unit) will be set by the tolerance of the ‘weakest link’, or the most antibiotic-sensitive community member. Alternatively, community tolerance may be higher than that of the weakest species in monoculture (the ‘community protection’ hypothesis), or lower (the ‘community sensitivity’ hypothesis). Higher than expected tolerance may occur if one or more species in a community excretes a compound which either actively degrades antibiotics in the medium (Perlin et al., 2009; Stiefel et al., 2015), or which activates tolerance mechanisms such as efflux pump expression in neighboring species (El-Halfawy and Valvano, 2012; Molina-Santiago et al., 2014). Lower than expected tolerance could result if sub-lethal concentrations of antibiotic, while not sufficient to arrest or kill any one species, sufficiently disrupt cross-feeding to inhibit community growth.

We tested the impact of cross-feeding on tolerance in a variety of conditions. We started with an engineered obligate mutualism involving *Escherichia coli, Salmonella enterica* serovar Typhimurium, and *Methylobacterium extorquens* (**Figure 1**, Harcombe et al, 2014). The tolerance of the mutualism was compared to the tolerance of each species in monoculture. These tests were done with ampicillin and tetracycline because the drugs have different mechanisms of action (Perlin et al., 2009, Hong et al., 2014), and because ampicillin can be enzymatically degraded (Munita and Arias, 2016), allowing the potential for cross-protection of sensitive species (Perlin et al., 2009). Further, we tested the impact of these antibiotics in both liquid media and agar plates to test the influence of spatial structure. Finally, to test the generality of our findings we investigated the effect of cross-feeding on tolerance in a model relevant for cystic fibrosis. This system involves the pathogen *P. aeruginosa*, and a previously defined anaerobic consortium of bacteria that provide carbon for the pathogen when growing on mucin (Flynn et al., 2016).

**Figure 1.**
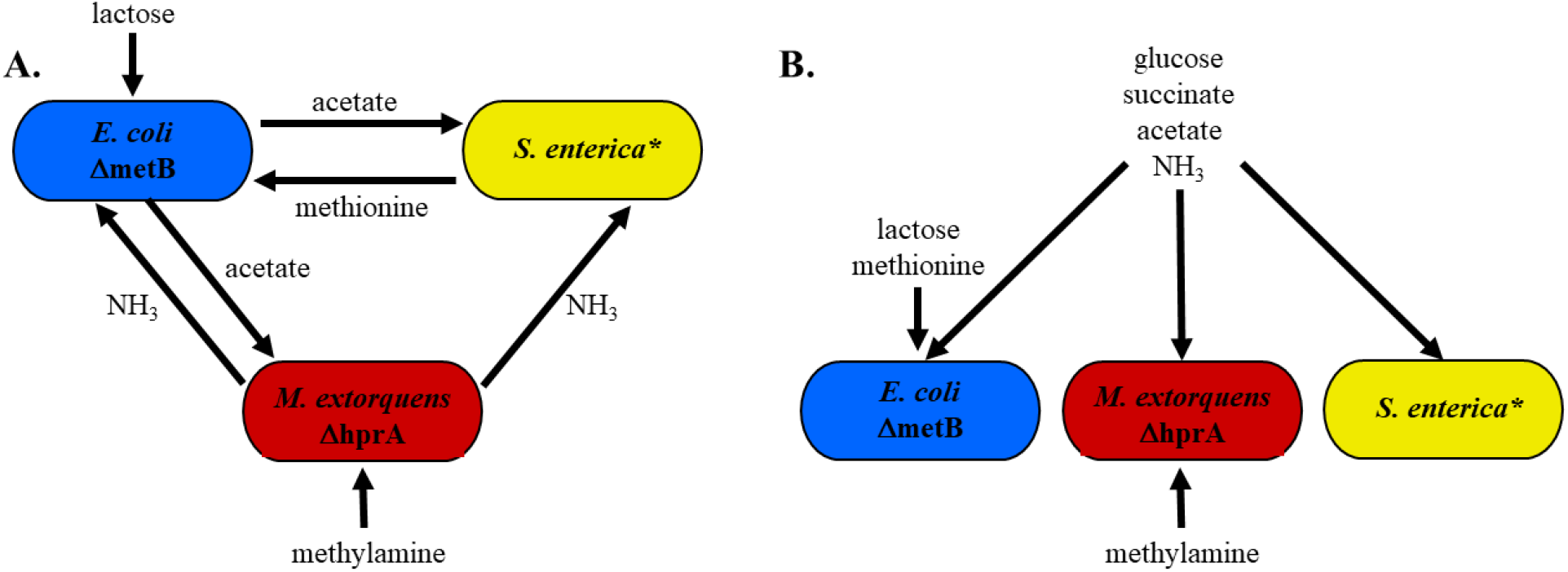
Cooperative and competitive model communities. **A.** Cooperative community. Methylamine and lactose are supplied in the growth medium as a nitrogen and a carbon source, respectively. *E. coli* consumes lactose and excretes acetate as a carbon source for *S. enterica* and *M. extorquens*. *S. enterica* secretes methionine for the methionine auxotroph *E. coli*. *M. extorquens*, which has a deletion in *hprA* which renders it unable to assimilate carbon from methylamine, provides nitrogen to the community via methylamine breakdown. **B.** Competitive community. Growth medium contains all metabolites necessary for growth of each individual species such that no cross-feeding is necessary to support growth.

Across all treatments and in both the model and clinically relevant systems, resistant bacteria were inhibited by lower concentrations of antibiotic when cross-feeding than when growing independently. However, we found that cross-feeding can provide protection to more sensitive bacteria. For both ampicillin and tetracycline, cases arose in which tolerance was higher than predicted based on measurements of tolerance in monoculture. Our results demonstrate that metabolic interactions impact antibiotic tolerance in a community, and suggest that antibiotic-resistant pathogens may be treated by targeting their less tolerant metabolic partners.

## Methods

### Bacterial strains and media

The three species community contained strains of *Escherichia coli, Salmonella enterica* and *Methylobacterium extorquens* all of which have been described previously (Harcombe et al., 2014). Briefly, the *E. coli* str. K12 contains a Δ*metB* mutation. The *S. enterica* mutant excretes methionine as a result of mutations in *metA* and *metJ* (Douglas et al., 2017, 2016; Harcombe, 2010). The *M. extorquens* AM1 Δ*hprA* mutant is unable to assimilate carbon from C1 compounds (Marx, 2008). In lactose minimal medium, the three species rely on each other for essential metabolites. *E. coli* secretes acetate byproducts which the other species rely on for a carbon source. *M. extorquens* releases ammonia byproducts which provide a source of nitrogen for other species. *S. enterica* secretes methionine, which is essential because our *E. coli* strain is auxotrophic for this amino acid. Each species has a unique fluorescent label integrated into its genome: cyan fluorescent protein (CFP) for *E. coli*, yellow fluorescent protein (YFP) for *S. enterica*, and red fluorescent protein (RFP) for *M. extorquens*. Bacteria were grown in minimal Hypho media containing varying amounts and types of carbon and nitrogen, depending on the medium type (see **Figure 1**). *Pseudomonas aeruginosa* strain PA14 (Rahme et al., 1995) was obtained from D.K. Newman (Caltech). The anaerobic consortium (composed primarily of *Prevotella sp., Veillonella sp., Fusobacterium sp., and Streptococcus sp*.) was derived from human saliva and was enriched using porcine gastric mucin as previously described (Flynn et al., 2016).

### Liquid media experiments

Bacteria were inoculated along an antibiotic gradient to measure the minimum inhibitory concentration (MIC) for monocultures and co-cultures. The OD of each culture was taken to ensure cells were in mid-log phase (OD~0.2-0.3) and cultures were diluted 1/200. Cells were inoculated into relevant wells on a 96-well plate, with fresh Hypho and varying concentrations of an antibiotic. The inoculate size for a species was kept constant at ~10^4^ in monoculture and community (i.e. community treatments started with 3x more total cells than monocultures). Ampicillin was used at 0, 0.2, 0.5, 1, 2, and 5 μg/mL for *E. coli, S. enterica*, and the communities, and at 0, 2, 5, 10, 20, and 50μg/mL for *M. extorquens*; these concentrations provided the best range of sublethal to lethal ampicillin concentrations. Tetracycline was used at 0, 1, 2, 3, 5, and 10μg/mL for *E. coli, M. extorquens*, and the community, and 0, 10, 20, 30, 50, and 100μg/mL were used for *S. enterica*. 96-well plates were placed into a Tecan InfinitePro 200 at 30°C for 120 hours. Measurements of OD600 were taken every 15 minutes to track overall bacterial growth, and fluorescence measurements were taken to track the growth of individual species. The minimum inhibitory concentration (MIC) was defined as the lowest antibiotic concentration at which no growth was seen by three times the lag time of the antibiotic-free control.

### Solid media antibiotic susceptibility experiments

Resistance on plates was determined by measuring the diameter of the zone of inhibition around an antibiotic containing disk. For monocultures, 150μL of log-phase culture (OD=0.5) was spread on Hypho plates (1% agar) in a lawn; for communities, 150μL of culture from each species was mixed, spun down, and re-suspended in 150μL of the appropriate community medium before plating onto Hypho plates. Discs of filter paper 6mm in diameter were inoculated with 25 μg antibiotic, and left to dry for 10 minutes. Discs were applied to the center of plates with bacteria, and plates were incubated at 30°C for 48 hours (*E. coli, S. enterica*, competitive community) or 72 hours (*M. extorquens*, cooperative community), depending on how long it took for cells outside the zone of inhibition to become confluent. The diameter of the zone of clearing was measured three times for each plate and averaged to provide a single plate measurement.

### Fluorescence microscopy

Fluorescent microscope images were obtained using a Nikon AZ100 Multizoom macroscope with a C1si Spectral confocal attachment, 4x objective lens at 3.40x magnification using Nikon NIS Elements software. The 457nm argon laser was used to visualize CFP, the 514nm laser was used for YFP, and the 561nm laser was used for RFP. Disc diffusion Petri plates were placed on the stage and images from 2x10 fields of view (for monocultures) or 2x12 fields of view (for communities) were obtained and stitched together using Elements software. Images for each fluorescent protein were overlaid using Fiji image analysis software (Schindelin et al., 2012).

### Testing β-lactamase production

Nitrocefin discs (Sigma-Aldrich, 49862) were used to determine if *M. extorquens* was producing a β-lactamase. For solid medium, cells were scraped off agar plates and suspended in the appropriate liquid medium; the OD600 of the suspension was then determined and diluted to an OD600 of ~0.5, to match the OD600 of liquid cultures. Discs were placed on a microscope slide and 15μL of liquid culture or diluted solid medium suspension was added to the disc. After 60 minutes, a color change from yellow to purple/pink indicated the production of a β-lactamase that hydrolysed the nitrocefin in the disc (O’Callaghan et al., 1972). As a positive control, an *E. coli* strain carrying a pBR322 plasmid, which contains a *bla* β-lactamase gene was also tested, and the plasmid-free *E. coli* strain was used as a negative control.

### *Pseudomonas aeruginosa* cross-feeding model

Antibiotic tolerance assays were performed in minimal medium containing 1 mM magnesium sulfate, 60 mM potassium phosphate (pH 7.4), 90 mM sodium chloride, trace minerals (Marsili et al., 2008) and supplemented with autoclaved and dialyzed pig gastric mucin (30g/L, Sigma-Aldrich) for co-cultures or glucose (12 mM) for *P. aeruginosa* monoculture. Ampicillin was added at indicated concentrations. For mucin fermenting community assays, cultures were inoculated from freezer stock characterized previously (Flynn et al., 2016) and allowed to grow under anaerobic conditions containing carbon dioxide, hydrogen and nitrogen (5:5:90) at 37 °C for 48 hrs. For *P. aeruginosa* assays, cultures were inoculated from overnight cultures grown in LB and allowed then to grow while shaking aerobically at 37°C for 16 hrs. Optical densities were determined using a Biotek Synergy H1 plate reader. Optical densities are given as mean and standard deviation of three replicates.

Cross-feeding assays were performed as described previously (Flynn et al., 2016). Briefly, a mucin fermenting community from freezer stock was inoculated into the minimal mucin medium described above and allowed to grow for 48 hrs anaerobically at 37°C. This culture was used to inoculate the lower phase (1:100 dilution) which contained 2 mL of minimal mucin medium, 1% agar, and supplemented with ampicillin as indicated. After solidification of the mucin fermenting agar cultures in 16mm glass culture tubes, *P. aeruginosa* PA14 was added to buffered media containing no mucin and 0.7% agar to 1/1000 from an LB overnight culture. This mixture was then added to the top of the mucin fermenting community. After agar solidification, the tubes were placed in 37°C. After 60 hrs, the top agar section (containing PA14) was removed, homogenized by pipette in sterile saline, serially diluted, and plated on Luria Broth agar plates to enumerate PA14 in each culture.

### Statistical analyses

For liquid and solid media assays, at least eight replicates of each treatment were obtained for each antibiotic. Pairwise comparisons between monocultures and co-cultures were conducted using a Mann-Whitney U test with an applied Bonferroni correction for ten multiple comparisons. For *P. aeruginosa* cross-feeding assays, triplicate experiments were performed for each antibiotic concentration and community type. Normalized CFU values were calculated by dividing CFU at each antibiotic concentration by the CFU at 0ug/mL ampicillin. Comparisons of normalized values at each concentration were performed using a Mann-Whitney U test.

## Results

### Dependence on other species reduces the amount of antibiotic necessary to inhibit resistant bacteria in liquid media

In monoculture experiments, one species was at least 25 times more resistant than others, though the most resistant species was different for each antibiotic. When grown in the presence of ampicillin, *M. extorquens* had a median MIC of 100 μg/ml, which was significantly higher than the 2 μg/ml and 1 μg/ml necessary to inhibit *E. coli* and *S. enterica* respectively (**Figure 2A**, *P*<0.0001 for each). For tetracycline, *S. enterica* had a median MIC of 50 μg/ml (**Figure 2B**). This was significantly higher than the median MIC of 5μg/ml in *M. extorquens* (*P* <0.0001) and 2 μg/ml in *E. coli* (*P* <0.0001).

**Figure 2.**
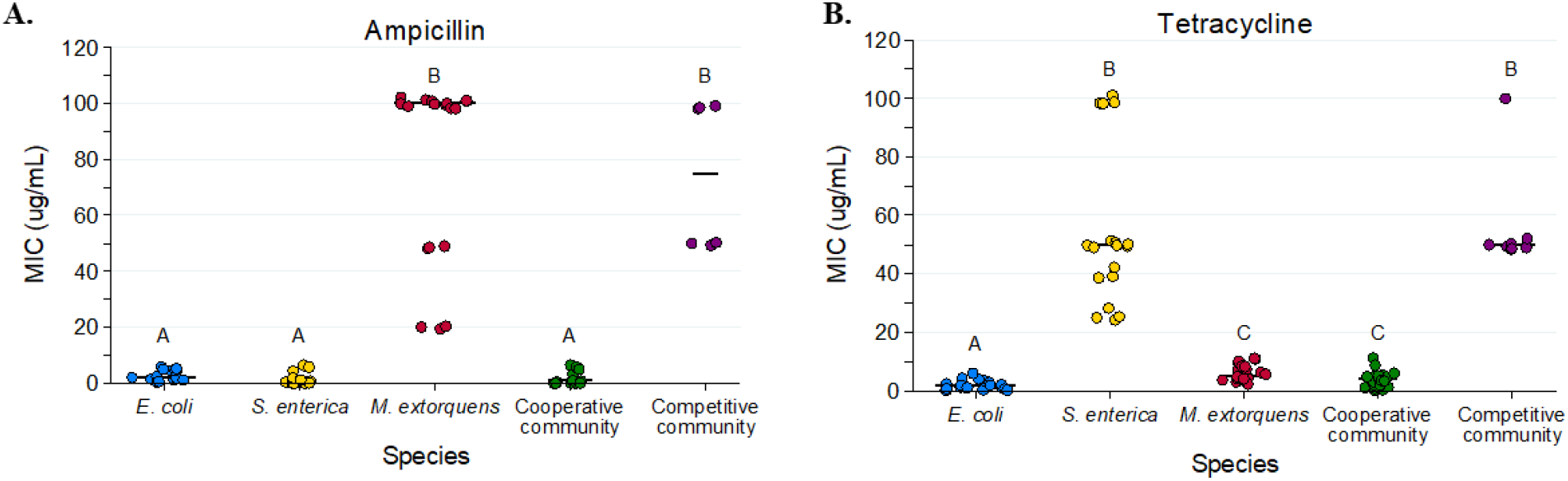
Minimum inhibitory concentration (MIC) values for each monoculture and community type in ampicillin (**A**) and tetracycline (**B**) based on OD600. Bars represent median values of MIC. At least eight replicates were performed for each species/antibiotic combination. Pairwise comparisons of median MIC were performed using a Mann-Whitney U test, with a Bonferroni correction applied for ten pairwise multiple comparisons. Letters above data points indicate significant differences between groups.

When bacteria were grown in an obligate mutualism rather than in monocultures, the amount of antibiotic needed to inhibit resistant species decreased substantially. Fifty-fold less ampicillin was needed to inhibit the cooperative community (1μg/mL) than to inhibit *M. extorquens* in monoculture (**Figure 3A**, *P* < 0.0001). Similarly, the median MIC of tetracycline for *S. enterica* decreased significantly from 50 μg/ml in monoculture to 4 μg/ml in the cooperative community (**Figure 3B**, *P* <0.0001).

**Figure 3.**
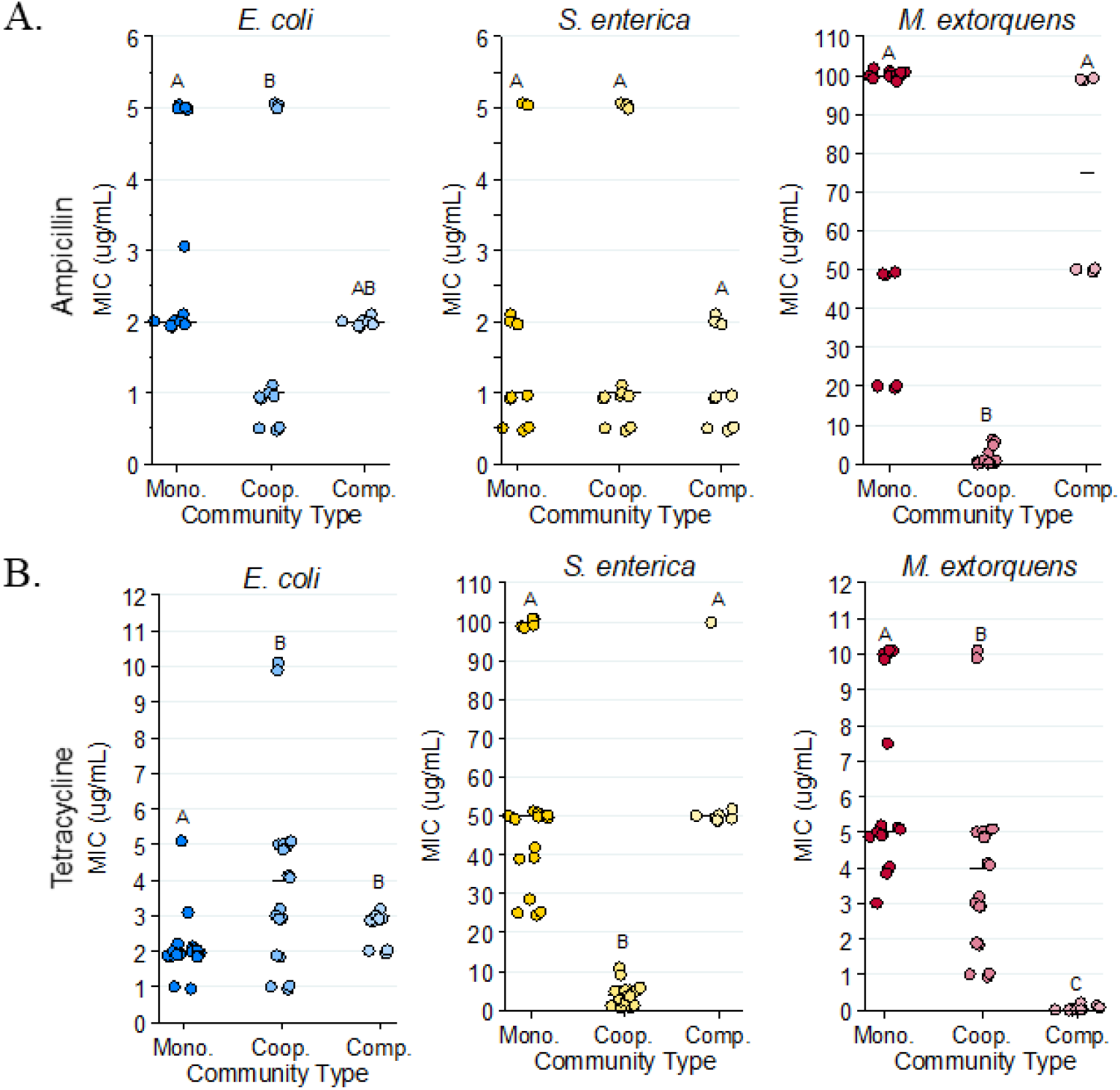
Minimum inhibitory concentration (MIC) values for each species in monoculture, cooperative community, and competitive community in ampicillin (**A**) and tetracycline (**B**). MICs were calculated based on fluorescence. Pairwise comparisons of median MIC were performed using a Mann-Whitney U test, with a Bonferroni correction applied for three multiple comparisons. Letters above data points indicate significant differences between groups.

We next distinguished the effect of species interactions from community complexity by measuring the MIC of the bacterial community when the species were not reliant on one another for resources (**Figure 1B, Figure 2**). The median MIC for resistant bacteria was not significantly different in the competitive community than it was in monoculture (for *M. extorquens* in ampicillin **Figure 3A**, *P* >0.90; for *S. enterica* in tetracycline, **Figure 3B**, *P*=0.80). Therefore, the decreased tolerance of resistant species to these antibiotics was a result of the metabolic interdependence, rather than simply the presence of other species.

We also repeated monoculture and competitive community experiments using acetate as the sole carbon source for *S. enterica* and *M. extorquens* to ensure that the change in carbon source between treatments was not responsible for changes in tolerance (**Supplementary figure 1**). We found that the same qualitative patterns were observed with *M. extorquens* showing high tolerance to ampicillin and *S. enterica* demonstrating high tolerance to tetracycline in both monoculture and competitive community. However, we found that growth was less robust, and that high initial concentrations of acetate may be toxic to *M. extorquens*.

### Tolerance of tetracycline was higher than expected in the cross-feeding community

Unexpectedly, the MIC of tetracycline for the cooperative community was higher than the least resistant monoculture, *E. coli* (median monoculture MIC 2μg/mL, median cooperative community MIC 4μg/mL, *P* =0.0111). This effect was not due to higher starting cell density in community (**Supplementary fig. 3A**). We therefore suspected the increased tolerance of *E. coli* in cooperative community might be due to increased lag times for *E. coli* in the cooperative community. As tetracycline is known to rapidly break down, we predicted that the slower growth in community might be protecting *E. coli* by allowing time for tetracycline to break down before growth started. Consistent with this prediction, the time to detectable *E. coli* growth was significantly longer in cooperative community than it was in monoculture (*P* = 0.003), and the MIC of *E. coli* in monoculture increased if media containing tetracycline was allowed to sit for 20 hours before cells were added (**Supplementary figure 3B**). Therefore, we concluded that the increased tolerance of *E. coli* to tetracycline in cooperative community was likely influenced by delayed *E. coli* growth relative to monoculture.

The competitive community appears to decrease the effective resistance of *M. extorquens* to tetracycline. In monoculture, the tetracycline MIC for *M. extorquens* was 5μg/ml, while in the competitive community *M. extorquens* growth was not observed even in the absence of antibiotic (**Figure 3B**). This indicates that competitive exclusion, rather than antibiotic tolerance, governs the growth of *M. extorquens* in this context. *M. extorquens* was not competitively excluded in the ampicillin treatments because it can grow at high ampicillin concentrations where its better competitors cannot. (**Supplementary figure 5**). This suggests that growth patterns of *M. extorquens* in tetracycline are governed by competitive ability rather than resistance, while in ampicillin *M. extorquens* experiences competitive release.

### Resistant bacteria are also constrained by sensitive partners in structured environments

We next tested whether growth on agar plates (rather than growth in liquid media) altered the impact that species interactions had on antibiotic resistance. We grew confluent bacterial lawns and measured the zone of clearing around an antibiotic disc at the center. All discs contained 2.5μL of 10mg/mL antibiotic. We hypothesized that spatial structure might enhance the ability of resistant bacteria to protect metabolic partners from degradable antibiotics like ampicillin, and thereby eliminate the reduction of tolerance in the cross-feeding system.

Resistance on agar largely mirrored results from the liquid media. As in liquid, *M. extorquens* was the most resistant monoculture on ampicillin, and *S. enterica* was the most resistant monoculture in tetracycline (**Figure 4**). Note that small clearing diameters signify high resistance, so the relative rankings in **Figure 4** are the inverse of **Figure 3**. We also once again observed that less antibiotic was needed to inhibit resistant bacteria in cooperative community than was needed in monoculture. Ampicillin cleared growth of *M. extorquens* out to a diameter of 32.7 mm in cooperative community, but only out to 15.7 mm in monoculture (**Figure 4A**, *P* = 0.002). Similarly, on tetracycline, *S. enterica* had a median diameter of clearing of 9.92 mm in monoculture and 33.7 mm in the cooperative community (**Figure 4B**, *P* <0.0001). In the competitive community, the zones of clearing matched those of the most resistant monoculture. Using acetate as the carbon source for the competitive community (in addition to lactose) and *S. enterica* and *M. extorquens* monocultures showed qualitatively similar results (**Supplementary figure 2**).

**Figure 4.**
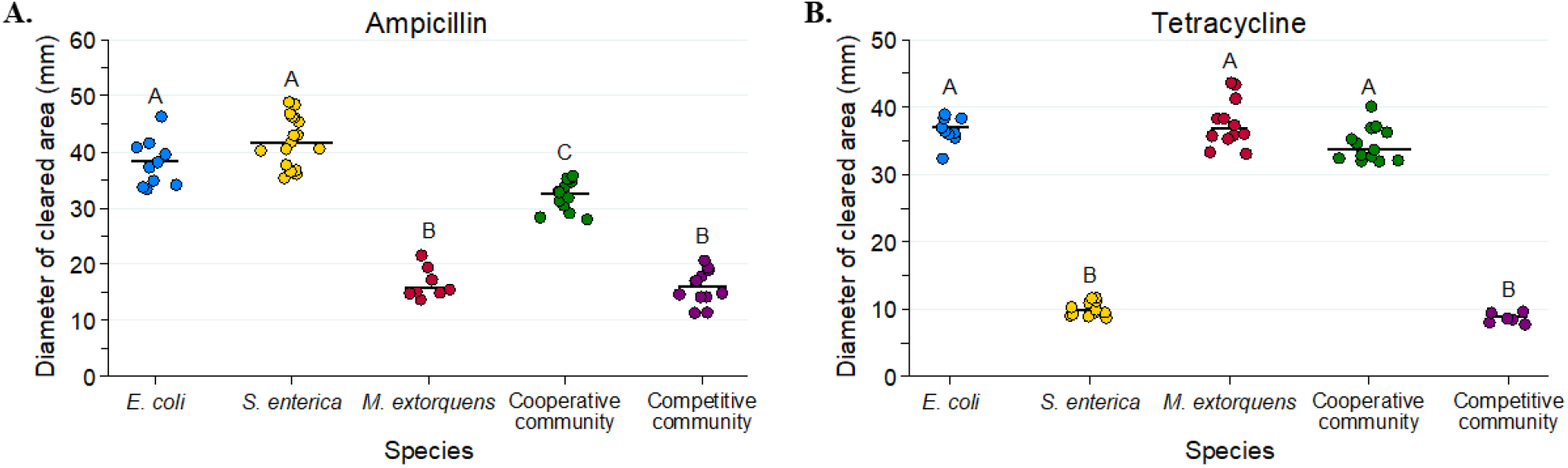
Diameters of zones of clearing for ampicillin (**A**) and tetracycline (**B**) disc diffusion assays. Pairwise comparisons of the zone of clearing for each monoculture and community was performed with a Mann-Whitney U test with Bonferroni adjustment for multiple comparisons. Significant differences are noted by different letters above each bar.

Though *M. extorquens* had lower tolerance for ampicillin in cooperative community, we did observe cross-protection of the more sensitive species. The cooperative community had a significantly smaller zone of inhibition than either *S. enterica* or *E. coli* monocultures (**Figure 4A**, *P* <0.0001 for *S. enterica* and *P* =0.01 for *E. coli*). Fluorescent images indicate that inhibition of both sensitive species was reduced in the presence of *M. extorquens* (**Figure 5**). In both cooperative (**Figure 5D**) and competitive (**Figure 5E**) community, *E. coli* and *S. enterica* grew closer to the disk of ampicillin in co-culture than they did in monoculture (**Figure 5A,B**). We found that protection was not due to an increase in initial cell density on community verses monoculture plates (**Supplementary figure 6A**), and that *M. extorquens* was responsible for providing the protection (**Supplementary figure 5B**). Consistent with the observed cross-protection, ampicillin degrading β-lactamases were found in the genome of *M. extorquens*. Further, nitrocefin disks were used to demonstrate β-lactamase activity when *M. extorquens* was grown on agar in the presence of ampicillin **(Supplementary Figure 7**).

**Figure 5.**
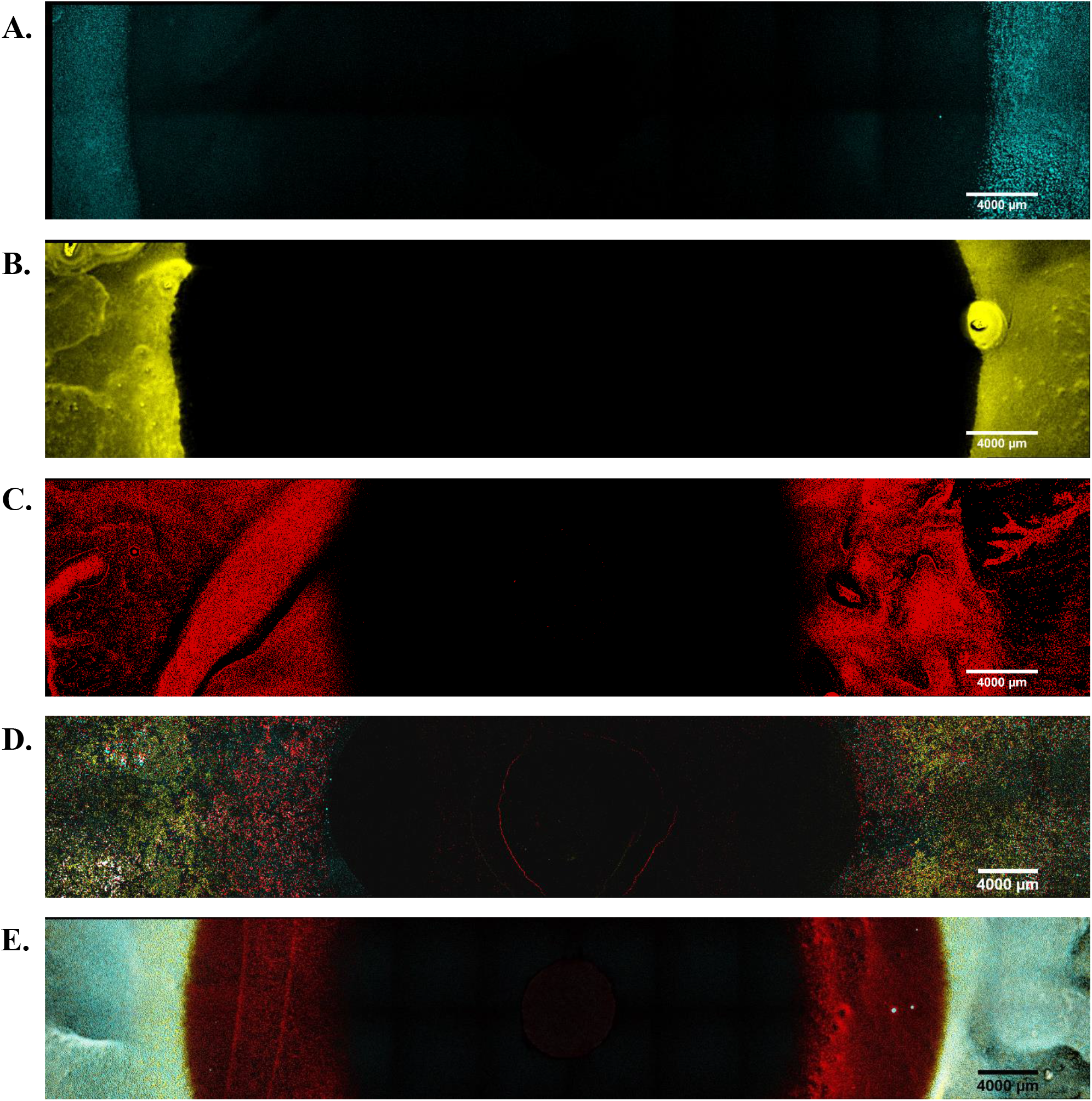
Fluorescent microscopy images of Petri plates with ampicillin antibiotic discs. *E. coli* fluoresces blue (CFP), *S. enterica* fluoresces yellow (YFP) and *M. extorquens* fluoresces red (RFP). (**A**) *E. coli* monoculture (**B**) *S. enterica* monoculture (**C**) *M. extorquens* monoculture (**D**) cooperative community (**E**) competitive community.

**Figure 6.**
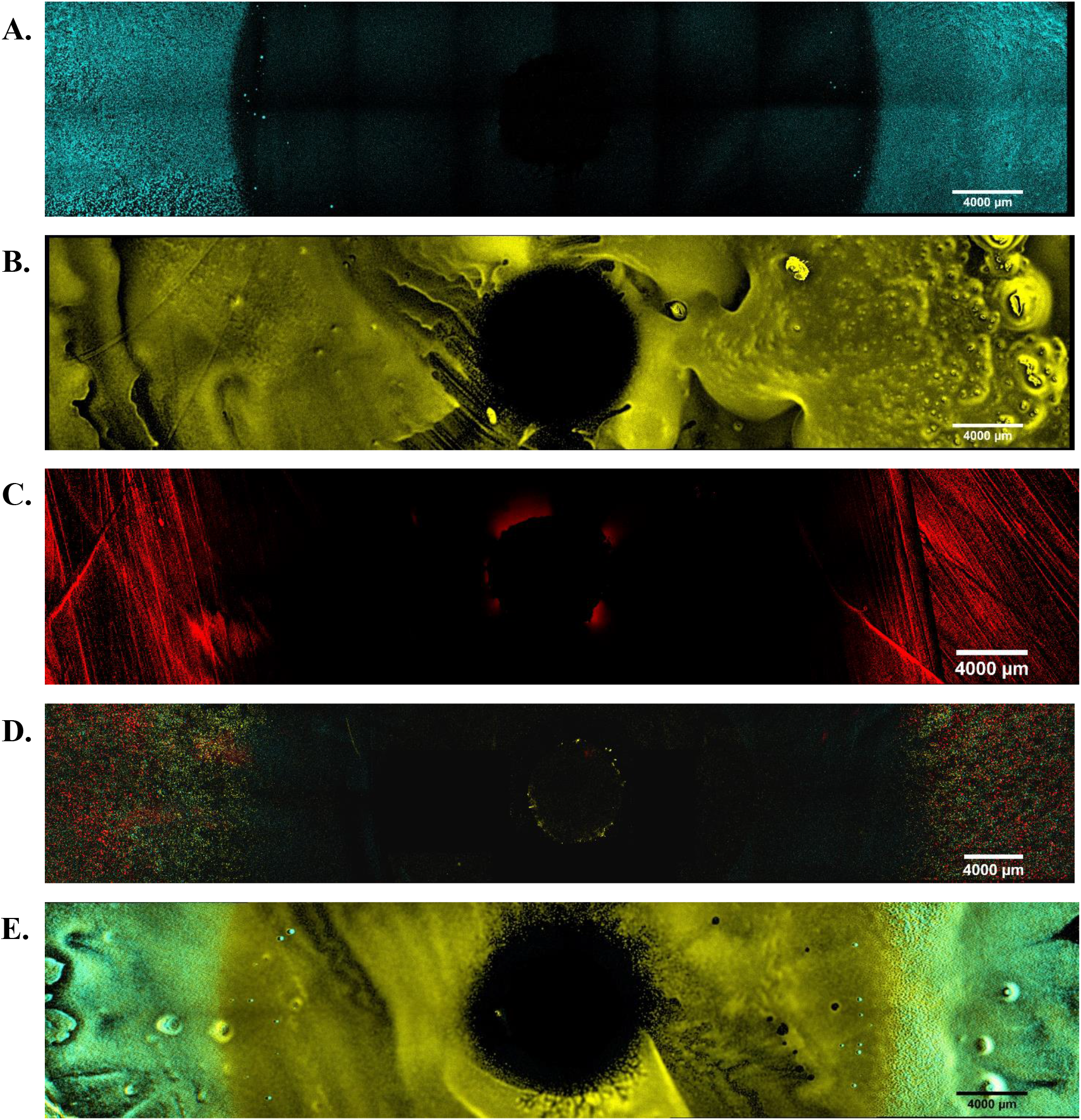
Fluorescent microscopy images of Petri plates with tetracycline antibiotic discs. *E. coli* fluoresces blue (CFP), *S. enterica* fluoresces yellow (YFP) and *M. extorquens* fluoresces red (RFP). (**A**) *E. coli* monoculture (**B**) *S. enterica* monoculture (**C**) *M. extorquens* monoculture (**D**) cooperative community (**E**) competitive community. No RFP signal was observed for the competitive community with a tetracycline disc.

### Metabolic dependency reduces antibiotic tolerance of a pathogen

To test whether our findings extend to medically relevant systems, we investigated how co-culturing influences the effective tolerance of *Pseudomonas aeruginosa* to ampicillin*. P. aeruginosa* is commonly found in the lung and is a leading cause of morbidity and mortality in people with cystic fibrosis (Langton Hewer and Smyth, 2017). Further, it was recently demonstrated that *P. aeruginosa* can cross-feed on carbon generated by mucin-degrading anaerobes (Flynn et al., 2016). In addition to its medical relevance, this system is distinct from our previous system in that the cross-feeding is not obligate (*P. aeruginosa* growth on mucin decreases but is not abolished by the absence of the anaerobes). We tested how ampicillin influenced the growth of *P. aeruginosa*, when grown alone on glucose versus in a cross-feeding co-culture.

Consistent with previous findings (Krogfelt et al., 2000), *P. aeruginosa* was highly resistant to ampicillin in monoculture (**Figure 7A**). No observable decrease in final yield was observed even out to 25 μg/ml of the drug. In contrast, the mucin-degrading anaerobes were inhibited by 5 μg/ml of ampicillin. Consistent with expectations, 5μg/mL ampicillin was sufficient to significantly reduce the density of *P. aeruginosa* grown in co-cultures on mucin, as compared to *P. aeruginosa* growth alone (*P* =0.0334). Similar results were observed if the *P. aeruginosa* monoculture was grown on mucin rather than glucose, however the maximum growth was substantially lower (**Supplementary Figure 8**). These data suggest that applying antibiotic to inhibit the growth of cross-feeding partners can inhibit resistant species even in a non-obligate cross-feeding system of clinical significance.

**Figure 7.**
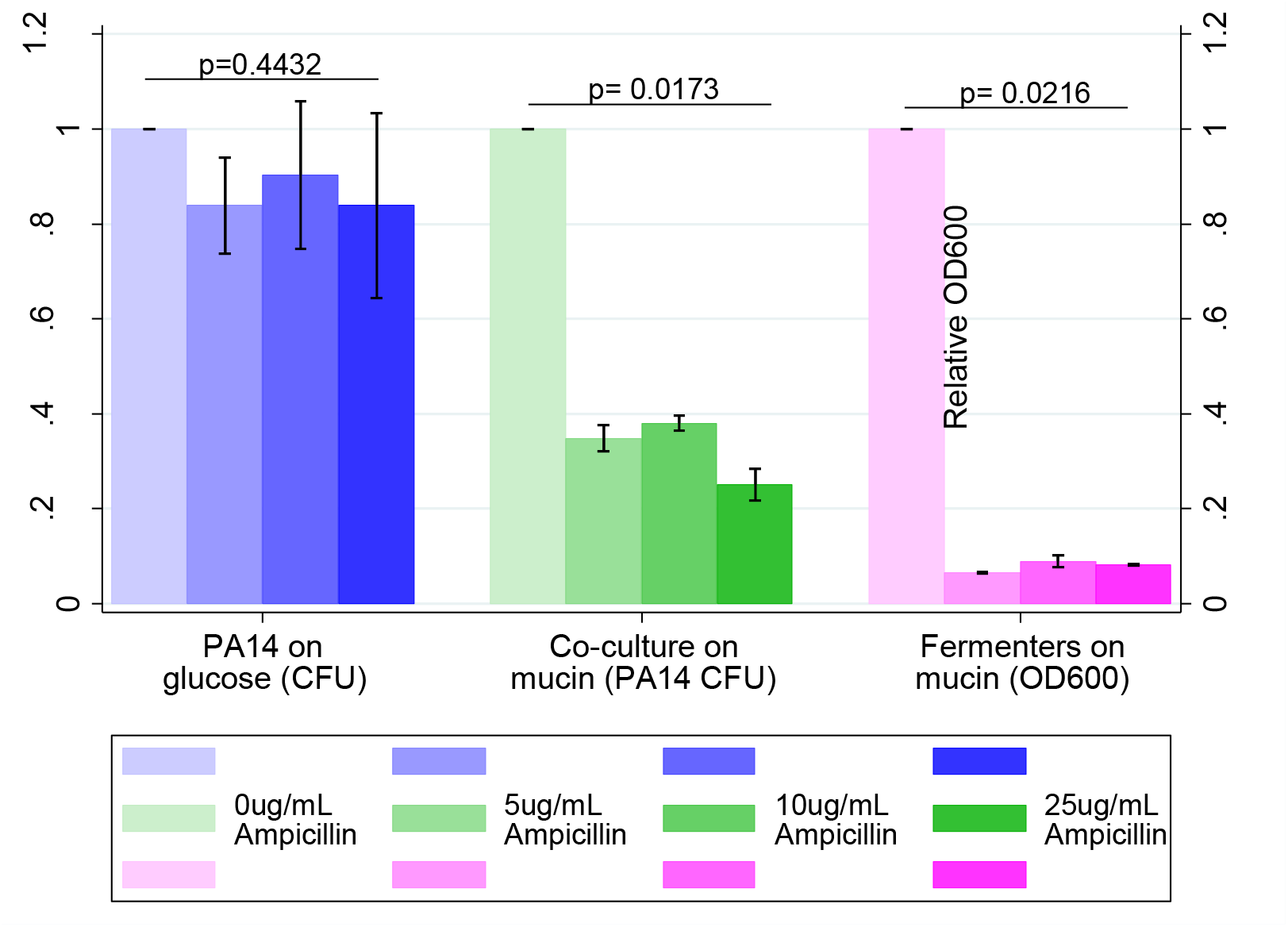
*Pseudomonas aeruginosa* PA14 growth alone verses cross-feeding with a mucin-fermenting community across a range of ampicillin concentrations. Growth of PA14 was measued by colony forming units (CFU). Growth of fermenters was measued by OD600. In all cases growth was standardized against the relevant antibiotic-free control. Each point represents the mean and standard deviation of triplicate samples. P-values were calculated using a Kruskal-Wallis test across ampicillin concentrations using relative CFU (for PA14) or OD600 (for fermenters).

## Discussion

Our results demonstrate that metabolic dependency between microbial community members plays a critical role in mediating the effect of antibiotics. In both mass-action (liquid) and structured (solid) environments we observed that bacterial species which show high levels of tolerance to a given antibiotic in monoculture are inhibited at much lower concentrations in an obligate mutualism. This was true for two antibiotics with different modes of action. Further, the effect of metabolic dependence on antibiotic tolerance extended to a medically relevant system with facultative cross-feeding. *P. aeruginosa* growth was reduced by substantially lower concentrations of ampicillin when the pathogen was cross-feeding off of mucin-degrading anaerobes that were sensitive to the drug. The constraint that cross-feeding places on the tolerance of bacteria was consistent across drugs and microbial systems.

We have shown that the ability of a given bacterial species to grow in the presence of an antibiotic is a combination of its intrinsic tolerance and the tolerance of species on which it relies for metabolites. Dependence on other bacteria reduced the MIC of bacteria with high resistance in monoculture. This change in MIC was driven by inhibition of a beneficial partner rather than a change in the resistance of the focal species. The effective tolerance of a cross-feeding network, therefore, is generally set by the ‘weakest link’ species; that is, the species with the lowest resistance to the antibiotic, whose tolerance in community usually matches its monoculture tolerance This suggests that antibiotics will often be more effective at controlling polymicrobial communities where there is extensive metabolic interdependence, and that tolerance in cross-feeding communities can be approximated from monoculture studies.

Unexpectedly, we did see deviations from our weakest link hypothesis. *E. coli* had a higher tetracycline MIC in the cross–feeding community than in monoculture, suggesting that cross-feeding provided some protection to *E. coli*. This slight, but significant, increase was likely driven by an increase in time to detectable *E. coli* growth when cross-feeding. Tetracycline breaks down rapidly, so delaying growth likely allowed microbial populations to experience reduced antibiotic concentrations. Similar antibiotic dynamics may often occur in clinical or environmental settings, (Yılmaz and Özcengiz, 2017, Oka et al., 1989), where metabolically inactive “persisters” commonly survive antibiotic treatment by delaying growth (Brauner et al., 2016). Community context further altered tolerance by enabling cross-protection of less tolerant species by more tolerant partners. On agar plates with ampicillin, both *E. coli* and *S. enterica* were able to grow closer to the antibiotic disk in the presence of *M. extorquens*. This protection was likely caused by degradation of the antibiotic as a result of β-lactamase activity in *M. extorquens*. Our results are consistent with previous observations that spatial structure can allow bacteria to lower local antibiotic gradients sufficiently to permit growth of sensitive isolates (Perlin et al., 2009). It is important to note, however, that there were limits on the extent of cross-protection in our community. The cross-feeding community increased the tolerance of *E. coli* and *S. enterica* but, the tolerance of *M. extorquens* was still lower in the cooperative community than it was in monoculture. Cross-protection may reduce the magnitude of the constraints placed on resistant species by their more sensitive metabolic partners, but it does not eliminate this constraint.

This study also demonstrates some issues which can arise when measuring the effect of antibiotics in microbial communities. It has previously been shown that MIC is a troublesome metric that can be influenced by factors such as changes in initial microbial density, or metabolic state (Artemova et al., 2015; Brauner et al., 2016). In our study, it was not possible to measure an MIC of tetracycline for *M. extorquens* in the competitive community, as *M. extorquens* was always outcompeted. The competitive release of *M. extorquens* in ampicillin treated competitive communities again deviates from standard patterns for MIC. Our results highlight that the community context further complicates the challenges associated with interpreting MIC measurements, especially for polymicrobial communities.

The constraint that cross-feeding places on antibiotic tolerance also extended to a polymicrobial community relevant to cystic fibrosis. It should be noted that this system involved facultative cross-feeding, so inhibiting anaerobes only reduced the yield of *P. aeruginosa* by two-fold. While this reduction is substantially smaller than the complete elimination of growth in the obligate system, two-fold changes may be medically relevant (Brotons et al., 2016; Cox et al., 2016; Pan et al., 2016; Priest et al., 2017; Weber et al., 2010). More broadly, the constraint in this treatment speaks to the generality of our findings. Even in scenarios with less extreme metabolic dependency, the impact of antibiotics can be magnified when highly resistant species are cross-feeding from less resistant species. Given that metabolic interactions are common in infection contexts (Murray et al., 2014; Ramsey et al., 2011), this work suggests that even narrow-spectrum antibiotics, designed to target a single species, may have ripple effects throughout a metabolically interconnected community.

Our results highlight that mutualistic networks are highly susceptible to environmental change. This result is consistent with work in other ecological systems from plant-pollinator to insect-symbiont (Gehring et al., 2014; Kikuchi et al., 2016; Silverstein et al., 2015). Integrating these ecological concepts into a microbial perspective may allow greater precision in our medical practices. Broad-spectrum reductions of bacteria in the gut can cause long-lasting negative health outcomes such as facilitating infections by *Clostridium difficile* (Chang et al., 2008). To develop precision treatments we need to be able to predict the impact of a drug on a focal population, and how a drug will affect off-target members of a microbial community. Our work highlights that precision microbiome management will require not only improved pharmacology but also a more comprehensive understanding of ecological interactions in microbial systems.

## Acknowledgements

We thank Michaela Muza for help with data collection. We also thank Ross Carlson for useful comments on the manuscript. This research was funded by Natural Sciences and Engineering Research Council of Canada and the National Institutes of Health.

